# Multi-level mosaicism shapes the evolution of small eukaryotic DNA viruses

**DOI:** 10.64898/2026.10.13.758929

**Authors:** Zachary K. Barth, Anh D. Ha, Frank O. Aylward

## Abstract

Eukaryote-infecting small dsDNA viruses, including those referred to as virophages, polintons, and their relatives, are major constituents of ecosystems across the globe as both agents of mortality and as integrated elements that shape host genome evolution. As the ubiquity of these viruses has become apparent, there have been efforts to organize their diversity into major lineages in order to better understand their biogeography, ecological impact, and evolution. Virophages, polintons, and polinton-like viruses were recently placed in the sub-phylum *Polisuviricotina,* though efforts to resolve relationships within this sub-phylum have been complicated by the vast diversity of these viruses and the existence of horizontal gene transfer between their genomes. To address these complications, we have used a gene profile occurrence based method of phylogenetic tree construction to aid in comparative genomic analysis of *Polisuviricotina*. We find that mosaicism is common within *Polisuviricotina*, in particular, reassociation of replication modules and morphogenic modules occurs throughout major *Polisuviricotina* groups. In addition to mosaic exchanges of full replication modules, we also observe extensive domain swapping within some replicases, indicating that mosaic recombination occurs on both the genome and subgenic level in these viruses. Analysis of individual replication modules suggests that they may be capable of some level of autonomous mobility, partially decoupling their evolution from the morphogenic module. This model helps explain the high levels of mosaicism in these viruses and has implications for the mechanisms through which small dsDNA viruses evolve.

## Introduction

Small dsDNA viruses of eukaryotes are increasingly recognized as major constituents of environments across the globe(Paez-Espino et al. 2019; C. M. Bellas and Sommaruga 2021; C. Bellas et al. 2023; Jeong et al. 2023; Chase et al. 2022; Starrett et al. 2021; Buck et al. 2024; Stephens et al. 2024; Piedade et al. 2024; Mimick et al. 2026). Central to both the history of this field are polintons (also called mavericks), named for their encoding of a protein primed PolB (pPolB) and a retrovirus-like integrase (rve-int), a type of DDE transposase (Kapitonov and Jurka 2006; Pritham et al. 2007). Polintons were initially thought to be self-synthesizing transposons before the discovery of double-jellyroll capsids (DJR) revealed that they represent endogenous viral elements (EVEs)(Krupovic et al. 2014). Subsequent discovery of virophages, small viruses that replicate in the presence of giant viruses in the group *Nucleocytoviricota*, uncovered another ubiquitous lineage of small dsDNA viruses(La Scola et al. 2008; Zhou et al. 2013). Some virophages were subsequently found to encode a pPolB and an rve-int, providing an intriguing link to polintons(Fischer and Suttle 2011; Koonin and Krupovic 2017). Later, a third group of similar viruses, polinton-like viruses (PLVs) was identified, which encode a DJR capsid and a variety of different replicases(Yutin, Shevchenko, et al. 2015). While at least one PLV has been shown to replicate autonomously(Pagarete et al. 2015), other characterized PLVs have been shown to replicate specifically in the presence of giant viruses(Stough et al. 2019; Thomas, Inamoto, Palermo, et al. 2026; Thomas, Inamoto, and Short 2026; Roitman et al. 2023), drawing another link to virophages.

In the last decade, a combination of metagenome sequencing and cultivation-based methods has uncovered a vast and rapidly expanding diversity of viruses related to polintons, virophages, and polinton-like viruses (all now included in the subphylum *Polisuviricotina, hereafter referred to as* polisuviruses)(Paez-Espino et al. 2019; C. M. Bellas and Sommaruga 2021; Chase et al. 2022; Piedade et al. 2024; Stephens et al. 2024; Mimick et al. 2026). Mining of eukaryotic genomes has revealed a remarkable diversity of these elements in diverse unicellular lineages(C. Bellas et al. 2023; Bulzu et al. 2025). The ubiquity of such viruses is intriguing given that some endogenous virophages restrict replication of their host virus(Koslová et al. 2024), and has led to the suggestion that these endogenous viruses may function as an inducible antiviral defense in protists in manner analogous to some bacteriophage satellite elements(Barth and Aylward 2024).

In addition to the many endogenous polisuviruses discovered through sequencing eukaryote genomes, metagenomic surveys from a variety of environments have recovered a broad diversity of these viruses; although many of these viruses are hyperparasitic entities or otherwise associated with giant viruses(La Scola et al. 2008; Fischer and Suttle 2011; Roitman et al. 2023; Koslová et al. 2024; Barth et al. 2024; Thomas, Inamoto, Palermo, et al. 2026), others may constitute autonomous viruses capable of infecting hosts independently(Pagarete et al. 2015). Collectively, the dramatic expansion of diversity of these small dsDNA viruses of eukaryotes has raised questions regarding their evolutionary links and the defining genomic features that could be used to demarcate certain lineages.

Understanding of the evolutionary relationships between polintons, virophages, and PLVs has been complicated by the multitude of common names that have been given to overlapping sets of viruses. For example, polintons and mavericks were discovered around the same time(Kapitonov and Jurka 2006; Pritham et al. 2007), and subsequent literature often refers to both names. Another group described viruses closely related to known polintons/mavericks and coined a third name, adintoviruses, highlighting the presence of an adenovirus-like cysteine-protease (adenain) within their genome(Starrett et al. 2021; Buck et al. 2024). In addition, recent work has been split over whether ‘PLVs’ should be understood as specific clade sister to polintons (Koonin, Fischer, et al. 2024), or if the label refers to a broader clade that would encompass the polintons as well(C. M. Bellas and Sommaruga 2021; C. Bellas et al. 2023). Lastly, the term ‘virophage’ has been used variously as a phylogenetic label or as a general descriptor for small dsDNA viruses that parasitize giant viruses(Roitman et al. 2023). To improve clarity in this field, there have been recent efforts to construct a formal taxonomy for these viruses. Most recently, polintons, PLVs, and virophages, have each been placed as separate classes in the subphylum *Polisuvirocotina* alongside an additional class composed of the vertebrate-infecting adenoviruses(Koonin, Fischer, et al. 2024). However, many unclassified viruses that likely belong to new classes likely remain to be demarcated, and the relationships of these lineages to classified groups in the *Polisuviricotina* remains unclear. For the purposes of this study, polintons can be understood as synonymous to members of the class *Polintoviricetes*, PLVs to *Aquintoviricetes*, and virophages to *Virophaviricetes*.

A major hurdle to understanding phylogenetic relationships between viruses is the prevalence of horizontal gene transfer and genome mosaicism within certain viral lineages. In broad terms, many ssDNA virus lineages appear to have chimeric origins that may have involved parallel acquisition of key replication and structural genes from common ancestral pools(Koonin et al. 2020) and a transition between different structural gene lineages is inferred to link the giant dsDNA nucleocytoviruses and mirusviruses(Gaïa et al. 2023). In addition to these major evolutionary transitions, fine scale genomic mosaicism is also known to occur, which has been most well-studied in tailed bacteriophages (class *Caudoviricetes*). The classic example of this is the dsDNA bacteriophage P22, which possess a lysogeny regulatory module with high similarity (up to 96% nt identity) to bacteriophage lambda, but has a short-tailed structural module unrelated to the long-tailed module of lambda(Ackermann 2015). In the laboratory, chimeras of morphologically distinct phages have arisen through illegitimate recombination(Ackermann 2015). While such chimeras are likely very rare in nature, bacteriophage metagenomic comparisons have revealed rampant recombination in the wild(Dion et al. 2020; Moura De Sousa et al. 2021). The dsDNA genomes of tailed phages show extensive turnover of accessory genes that defend against other phages or subvert host immunity, and gene conversion of essential genes into highly sequence-diverged homologs or unrelated functional analogs appears to be a common occurrence(Edgell et al. 2010; Pope et al. 2013; Barth et al. 2023). The genomic plasticity of bacteriophages as well as their vast diversity has made protein alignment based phylogenomic methods inadequate for determining the relationships between certain bacteriophage genomes. This has led some to adopt alternative methods for bacteriophage phylogenetics such as proteomic or gene profile occurrence tree construction(Rohwer and Edwards 2002; Nishimura et al. 2017; Meier-Kolthoff and Göker 2017; Aiewsakun et al. 2018; Low et al. 2019; Weinheimer et al. 2025).

Given the remarkable diversity of polisuviruses in the environment and abundance of several as-yet classified lineages in global metagenome datasets, we performed a large-scale comparative genomic analysis of various lineages of *Polisuviricotina* to shed light on the evolutionary relationships between these groups. We find that these method groups *Polisuviricotina* lineages primarily on the basis of their morphogenic module. These lineages exhibited unstable associations with multiple classes of replicases, suggesting extensive horizontal transfer of replication genes and allowing for the independent evolution of morphogenic and replication gene modules.

## Results and Discussion

### Gene content and phylogeny of *Polisuviricotina*

To better understand the diversity of *Polisuviricotina*, we constructed a set of orthologous groups (OGs) from the proteins encoded by a representative sample of sequenced *Polisuviricotina* genomes. We sampled viral genomes from a wide range of both cultivated and metagenome-derived datasets(Paez-Espino et al. 2019; C. M. Bellas and Sommaruga 2021; Stephens et al. 2024; Jeong et al. 2023; Koslová et al. 2024; Buck et al. 2024; Starrett et al. 2021; Yutin, Shevchenko, et al. 2015; Bao et al. 2015) as well as from EVEs reported in protist genomes(C. Bellas et al. 2023). The genome set included 1055 total genomes and included representatives from three formally-classified lineages (*Virophaviricetes*, *Polintoviricetes*, and *Aquintoviricetes*) as well as a number of unclassified elements. As expected, core functions were well represented in our OG set, with the DJR-fold major capsid protein (MCP), minor capsid (mCP), A32 packaging ATPase, and replicase being the four most abundant functional classes, with lipases and cysteine-proteases being fairly common as well (Figure 1.) Somewhat surprisingly, the fifth most common functional class was nucleases, which can have a wide range of functions in viral genomes. Although it is unclear what role these enzymes play, one possible role would include genomic defense against other viruses or host antiviral mechanisms. Similarly, methyltransferases and TET-dioxygenase coupled glycosyltransferases(Burke et al. 2021) were prominent among our data set, suggesting that these viruses may encode restriction modification (RM) systems. The presence of RM systems in polisuviruses might indicate similarities to integrative bacteriophage, which often encode anti-viral defense. Integrated *Polisuviricotina* genomes might provide similar defense for unicellular eukaryotes, which lack the complex structural defenses of plants(Doughari 2015) and the specialized immune cells of animals. Additionally, polisuviruses encoded many predicted membrane pore-forming proteins (Figure 1) with similarity to bacteriophage holins (OG032, OG081, and OG228, Table S1) as well SLATT domain encoding proteins (OG019, OG097, OG141, Table S1) which are common in antiviral systems encoded by bacteria. *Polisuviricotina* genomes encoded pore forming enzymes could conceivably be involved in either abortive infection defenses against other viruses, or lytic egress from the host cell. Overall, our results demonstrate that *Polisuviricotina* OGs are enriched for core viral functions, and also include a range of protein families that may facilitate conflicts with other viral genomes.

**Figure 1.**
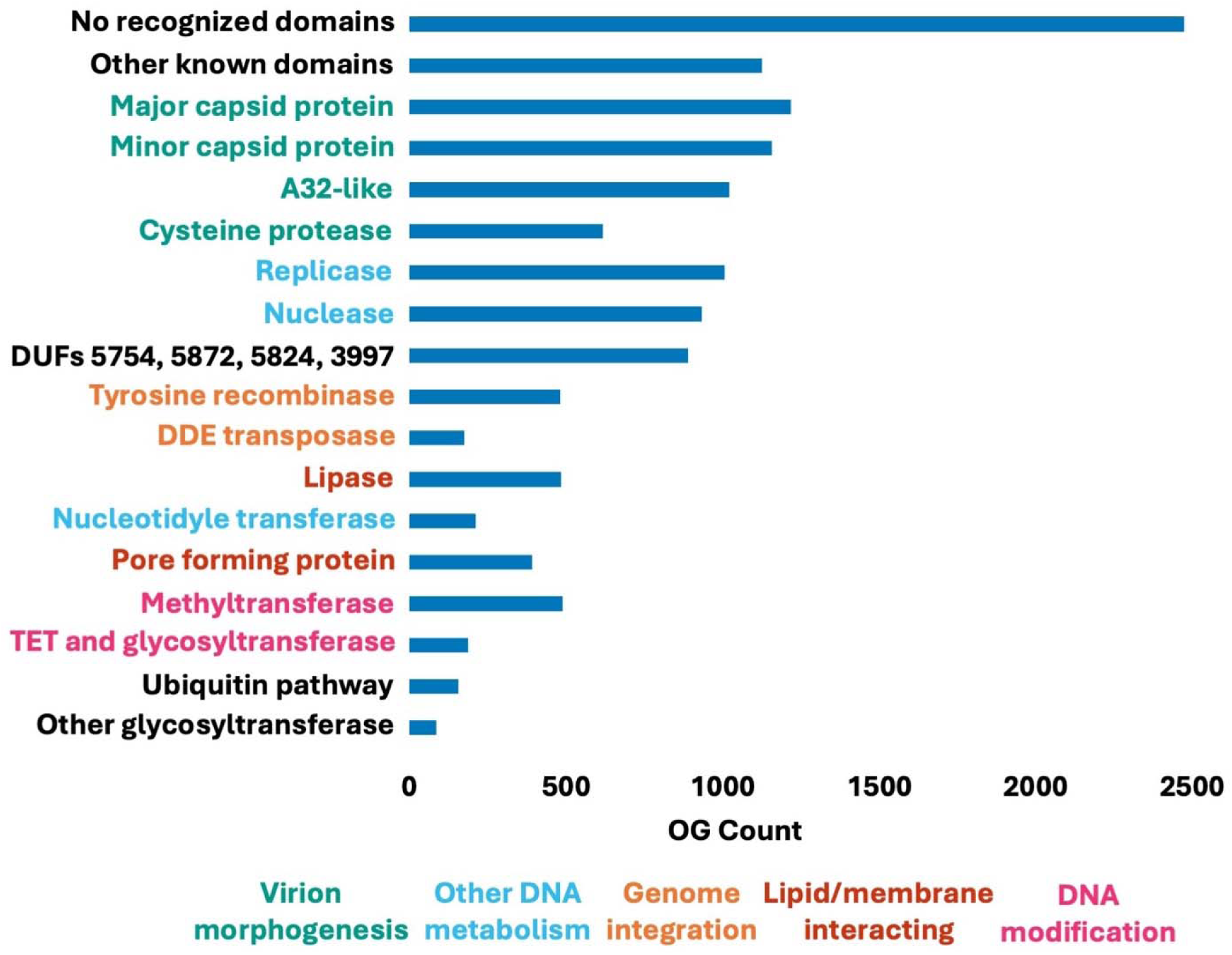
Polisuviruses possess diverse gene repertoires. Counts for the occurrence of functional categories of different OGs across our polisuvirus database. Each coding sequence within a genome is counted as one occurrence for the OG containing that sequence, and multiple OGs were collated into functional classes based on their annotations. Functional classes are colored by broader functional categories for genes involved in virion morphogenesis, DNA metabolism, genome integration, lipid and membrane interacting pathways, and DNA modification. The annotations for OGs and custom profiles can be found in Table S1.

We found a diversity of replication genes within polisuvirus genomes. Five OGs (OG021, OG038, OG055, OG115, and OG135) corresponded to pPolB proteins (Table S1), and additional curation split OG038 and OG135 into three and two profiles respectively. Six OGs (OG049, OG060, OG098, OG169, OG176, and OG183) corresponded to superfamily 1 (SF1) helicases (Table S1). Recent work has shown that the PLV encoded Tlr1-like SF1 helicases have a divergent PALM domain detectable through structural prediction(Koonin, Fischer, et al. 2024; Krupovic et al. 2024). The PALM domain is the catalytic domain of pPolB, suggesting that many PLV encoded SF1 helicases function as autonomous replicases(Koonin, Fischer, et al. 2024; Krupovic et al. 2024). We additionally found one Herpes Origin-binding protein-like (OBP) superfamily 2 helicase OG (OG052) and four D5-like superfamily 3 helicase OGs (OG000_M, OG002, OG203, and OG2011). Both OBP and D5-like helicases most often occurred alongside archaeal-eukaryote primase (AEP) or PolA domains (Data S1). We also detected one OG (OG200), that appeared to be related to a circovirus Rep-like nickase. These proteins consist of an HUH endonuclease domain and a superfamily 3 helicase. In circoviruses, these proteins initiate host polymerase mediated rolling circle replication of viral genomes(Tarasova et al. 2021). Similarly, a number of D5-like helicases occurred as stand-alone proteins suggesting that they may function in initiating replication by separate replication proteins encoded by polisuvirus, giant virus, or cellular genomes.

We next sought to construct a global phylogeny of polisuviruses so that we could better examine broad evolutionary trends in this group. Global phylogenetic analysis of polisuviruses is not possible using traditional marker gene approaches given the variable genetic content across groups together with the high levels of sequence divergence between different lineages, so we implemented a recently-developed approach that uses phylogenetic algorithms to generate a tree based on an occurrence profile of the OGs in each genome. This approach has recently successfully implemented for bacteriophages, which also have highly mosaic genomes that are highly divergent across lineages(Weinheimer et al. 2025). Within the tree we generated using this approach, viruses were clustered largely based on the occurrence of MCP OGs and produced major clades corresponding to the *Aquintoviricetes, Virophaviricetes,* and*Polintoviricetes,* demonstrating that this method is capable of reproducing established ICTV groupings (Figure 2.)

**Figure 2.**
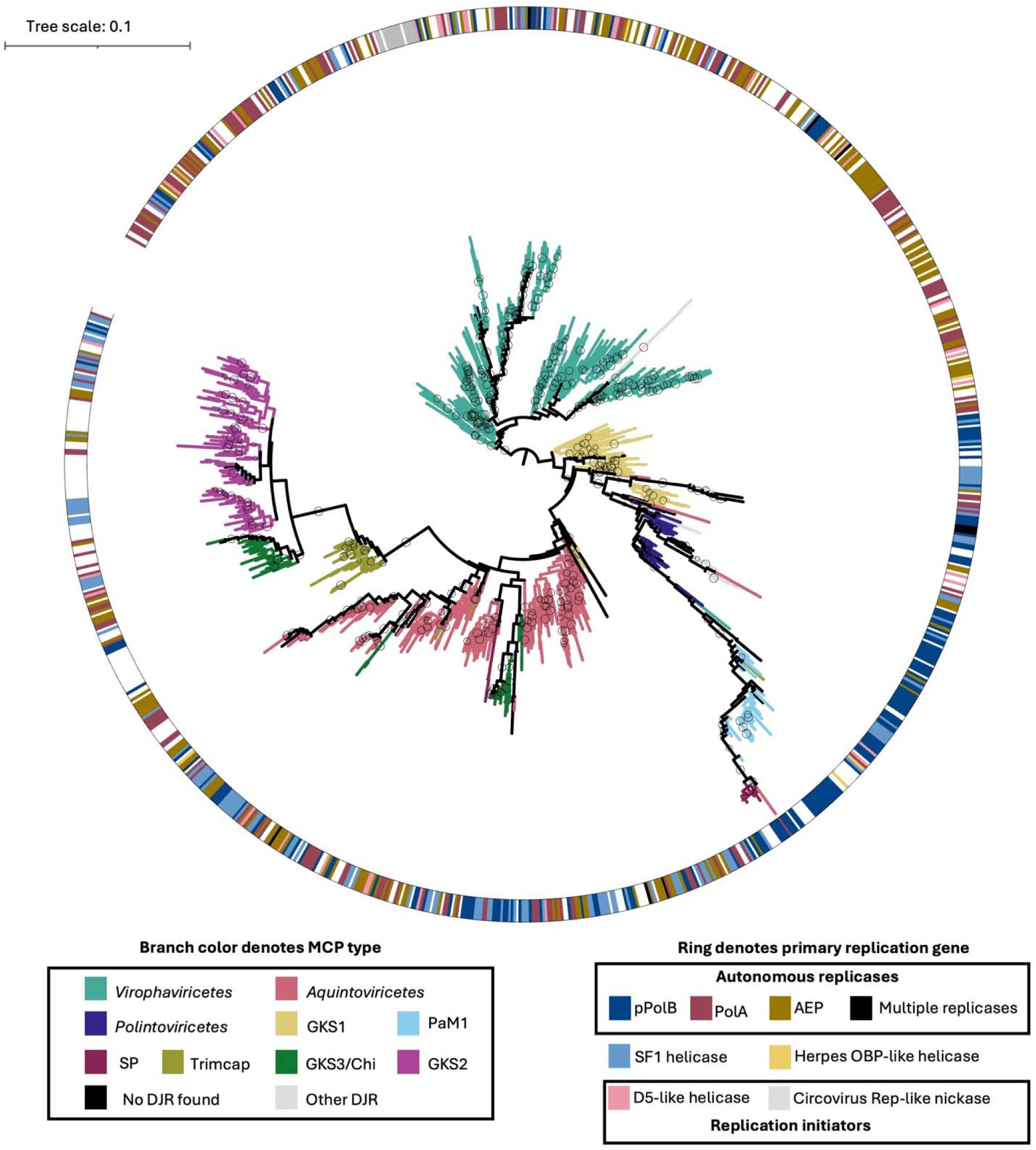
A trait based phylogeny recapitulates major classes of polisuviruses. An OG occurrence trait based tree for polisuviruses. Branches are colored according to MCP lineage, while the outer ring denotes replicase type. The presence of recognized replication genes is denoted by the outer ring. In cases where the genome encodes both a putative replicase and a replication initiator, only the replicase is indicated. The tree was artificially rooted at the base of the Virophaviricetes. Circles on branches represent bootstrap values greater than or equal to 90. The binary occurrence matrix used to generate this tree can be found in Data S1.

In addition to MCP profiles recognized as belonging to the three official polisuvirus classes, our approach also generated profiles equivalent to several MCP clusters that have been previously reported but have not yet been worked into the ICTV taxonomy. Notable these are the GKS1, GKS2, and GKS3-Chi MCP groups abundant in freshwater lakes and named for the Gossenköllesee lake in Austria(C. M. Bellas and Sommaruga 2021; C. Bellas et al. 2023), as well as MCPs from the Trimcap group which is belong to genomes abundant in marine environments that encode three separately evolved MCP genes(Piedade et al. 2024; Mimick et al. 2026; C. Bellas and Sommaruga 2026). While none of these viruses have been incorporated into ICTV taxonomy the name gosseviruses has been proposed for GKS1-type MCP encoding viruses, and the name alpenseeviruses has been proposed for GKS2-type MCP encoding genomes. Interestingly, the *Polintoviricetes* clade was nested within a larger clade comprised of gosseviruses and contained subclades defined by the PaM1-type MCP named for *Paulinella micropora* and abundant in *Rhizaria* and *Amoebazoa*(C. Bellas et al. 2023), as well as previously described Entomopoxvirus associated polisuviruses(Barth et al. 2024). This architecture raises the question of whether these groups should be considered part of *Polintoviricetes* as well or whether they comprise distinct lineages. Notably, recent work has grouped GKS1 MCPs with those of polintons(C. Bellas and Sommaruga 2026). Our tree also produced monophyletic clades for the Trimcap group and alpenseeviruses, suggesting that they may represent additional classes within *Polisuviricotina*. Interestingly, the GKS3-Chi MCP type was split between two separate clades, one sister to the alpenseeviruses, and another nested within the *Aquintoviricetes*.

Our phylogenetic approach does not show strong support for some aspects of the current organization of the *Virophaviricetes* below the class level (Figure S1). *Virophaviricetes* is currently organized into four orders: *Divpervirales* containing the rumen associated virophages, *Lavidavirales* containing Mavirus*, Mividavirales* containing virophage Sputnik, and *Priklausovirales* containing a number of viruses discovered through metagenomic sequencing(Roux et al. 2023). *Priklausovirales* contains four separate families: Burtonviroviridae, *Dishuiviroviridae*, *Gulliviroviridae*, and *Omnilimnoviroviridae*. Our workflow generated separate orthologous gene groups for the MCPs from *Lavidavirales*, *Mividavirales*, and *Gulliviroviridae*, as well as a fourth orthologous gene group that encompassed the remainder of *Priklausovirales* This notably splits *Gulliviroviridae* from the rest of the *Priklausovirales*, We did not observe a unique OG profile for the *Divpervirales* despite this order being known to have a divergent capsid(Roux et al. 2023). This may be because *Divpervirales* genomes are rare relative to those of the other orders. Of the three different virophage MCP orthologous groups we detected, our tree only produced a monophyletic clade for the *Lavidavirales* -type MCP, but the *Gulliviroviridae*-type and *Mividavirales-*type were dominant within two subclades (Figure S1).

We also examined the relationship between MCP and mCP orthologous gene groups (Figure 3A-C). Occurrence of the minor capsid protein appears to have a strong influence on the topology of our tree. Each of the major *Polisuviricotina* classes in our tree contained multiple orthologous gene groups for their MCPs with *Polintoviricetes* split into the Mavpol1 and Mavpol2 groups, *Aquintaviricetes* split into the TsV, PgVV, and Dino groups, and the *Virophaviricetes* split into the *Lavidavirales*, *Mividavirales*, *Gulliviroviridae*, and an additional MCP OG that includes the remained of *Priklausovirales* that have designated ‘Other Virophage,’ (Figure 3A). For each of these MCP OGs, there were shared associations between the major and minor capsid protein groups. The *Virophaviricetes* primarily paired with two minor capsid orthologous groups (OG005 and OG000_J), with the vast majority of *Virophaviricetes* genomes encoding the OG005 mCP(Figure 3A).

**Figure 3.**
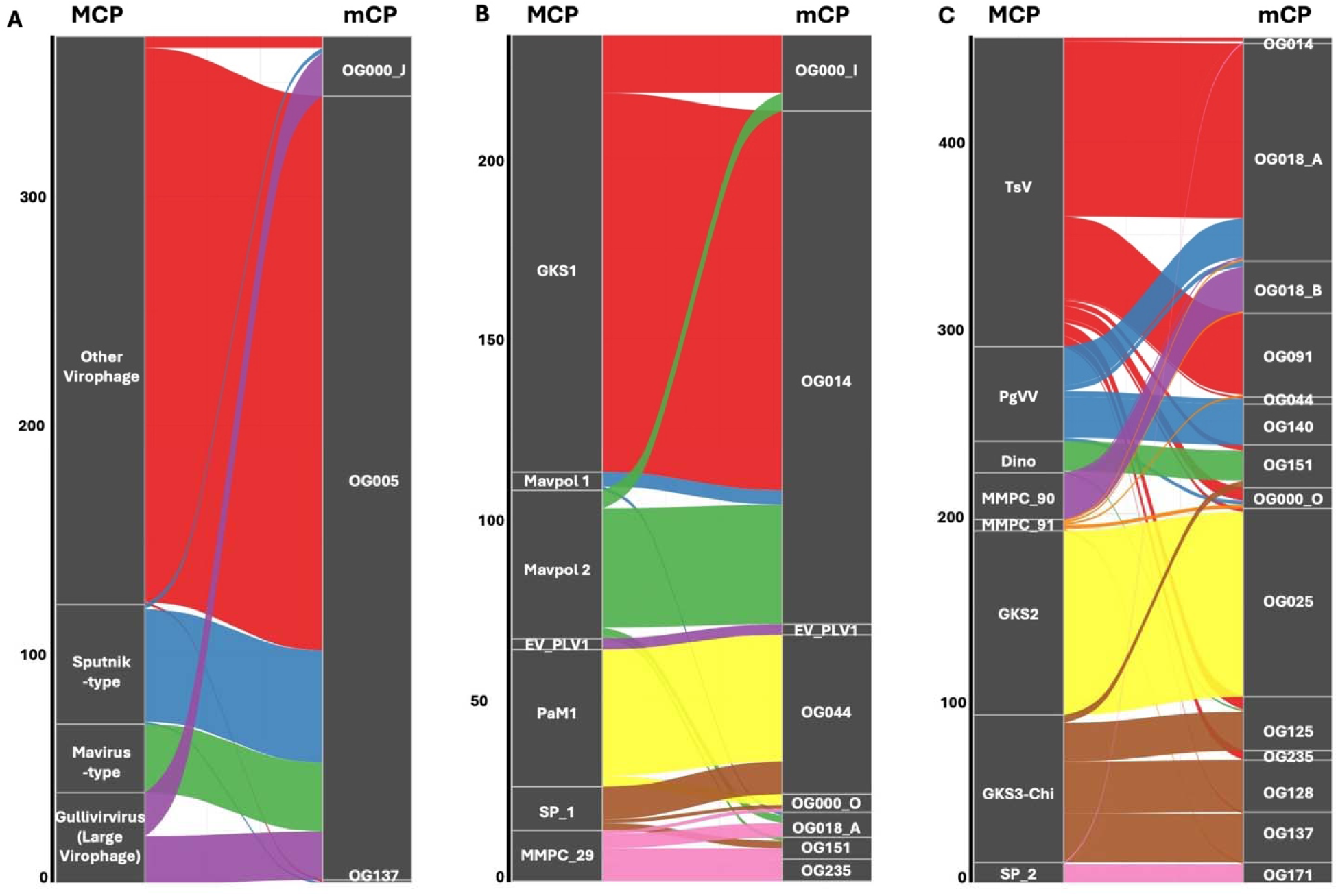
Major capsid protein and minor capsid protein OGs have distinct associations. Alluvial plots showing the associations between major capsid protein (MCP) and minor capsid protein (mCP) OGs in polisuviruses. Polisuviruses were organized into three putative clades based on results from figure 2: one group is comprised of Virophaviricetes (A); another group is comprised of gosseviruses, the Polintoviricetes, and genomes that placed within the Polontoviricetes (B); and the third group is comprised of the Aquintaviricetes, alpenseeviruses, Trimcap MCP encoding and GKS3-Chi type MCP encoding genomes. The number of genomes encoding the OGs is represented on the y-axis

Similarly, both Mavpol MCP groups paired primarily with the OG014 minor capsid group. The nesting of *Polintoviricetes* within the gosseviruses is likely because both have affinity for the OG014 and OG000_I mCP OGs (Figure 3B). A previous network analysis established that the three sets of MCPs for the *Polintoviricetes*, *Aquintaviricetes*, and *Virophaviricetes* are each related to each other(C. Bellas et al. 2023). While our phylogenetic approach risks dividing these sets into multiple traits, our results show that the phylogenetic affinity of these genes can still be captured through connectivity to other shared traits (in this case the minor capsid genes). Shared connectivity can also reveal previously unknown relationships, such as the link between *Polintoviricetes* and gosseviruses. This illustrates that the morphogenic module provides a strong phylogenetic signal through our approach.

### Polisuviruses exhibit hypermosaic genomes

There is some published evidence for mosaicism amongst the polisuviruses. A number of recognized ‘polintons’ including ones found in *Guillardia theta*, *Nematostella vectensis*, and *Trichomonas vaginalis*, possess major capsid proteins (MCPs) that do not belong to the maverick-polinton MCP clade(C. Bellas et al. 2023; Koonin, Fischer, et al. 2024), and a recently published evolutionary scenario for polisuviruses featured HGT of replicase classes within the virophage and PLV lineages(Koonin, Fischer, et al. 2024). Additionally, a group of entomopoxvirus associated polisuviruses exhibits signatures of genome mosaicism in their relatively recent evolutionary history(Barth et al. 2024). Genome mosaicism is a potential challenge for taxonomic organization of *Polisuviricotina*, as the PLVs (class *Aquintoviricetes*) and Virophages (class *Virophaviricetes*) are primarily defined by their MCPs, while the polintons (class *Polintoviricetes*) were initially defined based on the presence of a polymerase (pol-) and integrase (-int) combination.

Unlike capsid type, replicase class did not appear to cluster well with the binary tree structure (Figure 4A), suggesting that replicases have undergone widespread horizontal gene transfer over the course of *Polisuviricotina* evolution. However, some lineages did show greater congruence between MCP and replicase class, with *Polintonviricetes* showing affinity for pPolBs, and the *Gulliviroviridae* group showing affinity for Circovirus Rep-like nickases. (Figure 4A). While replicase class did not appear to correlate well with MCP type, there does appear to be an association between replicase class and integration genes (Figure 4B). There is a strong correlation between DDE transposases and PolB proteins as well as between tyrosine recombinases and PolA and AEP replicases. A functional link between protein primed PolBs and DDE transposases is possible due to their enzymatic activity. Integration modules typically act upon substrates located at the ends of the integrated genome(Hickman and Dyda 2015), and pPolB genes are typically primed by a terminal protein that attaches to the end of the viral genome(Salas 1991). As a result, pPolB and DDE proteins are likely to act upon the same genomic end sequence through either shared DNA specificity or through one protein acting as an adapter to provide DNA sequence specificity to the other protein. A functional link between PolA or AEP replicases and tyrosine recombinases appears more elusive. One possibility is that PolA and AEP replicases are able to elongate off of the cleaved recombination intermediates generated by the tyrosine recombinase. It should be noted that unlike DDE transposases, tyrosine recombinases typically generate circular excised products, though viral genomes are often linearized prior to virion packaging.

**Figure 4.**
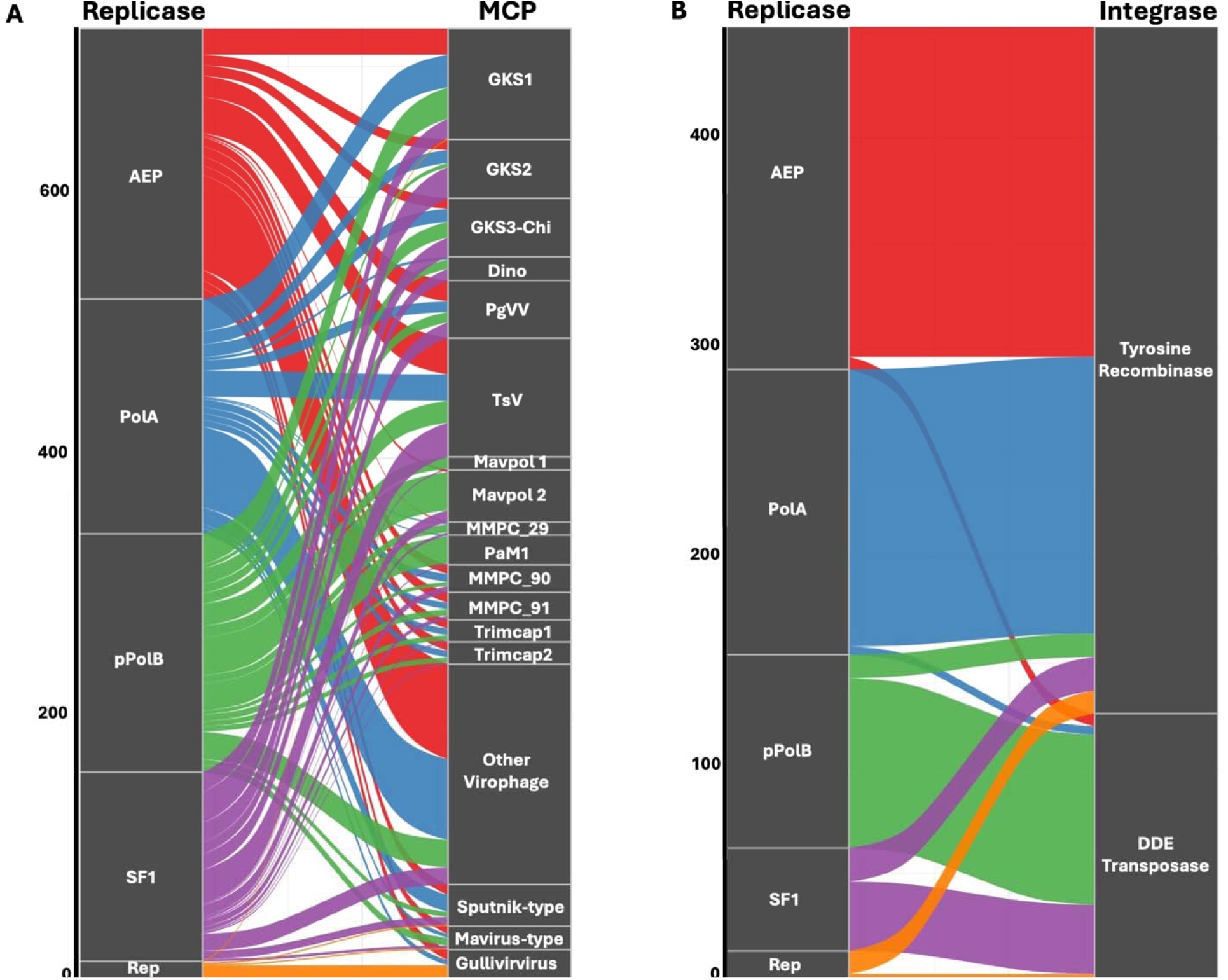
Replicases associate unstably with major capsid proteins and stably with integration genes. Alluvial plots showing the association between polisuvirus replicases and either major capsid proteins (MCP) (A) or integration genes (B). The number of genomes encoding the OGs is represented on the y-axis

To examine how the evolutionary history of polisuvirus pPolB clades is linked to morphogenetic modules, we constructed a tree of pPolBs and mapped their morphogenetic module groups on the outer ring (Figure 5A). Overall, the tree shows stable associations of pPolB and MCP types for some smaller subclades, but there is extensive mobility of pPolB types between different *Polisuviricotina* classes. Previous work has shown horizontal transfer of pPolB between the Polinton 1 and Polinton 2 subclades(Jeong et al. 2023), and the existence of a polinton-related pPolB in the ‘rumen associated virophage’(Yutin, Kapitonov, et al. 2015; Roux et al. 2023; Koonin, Fischer, et al. 2024). Our results suggest that these horizontal transfers may not be exceptions to *Polisuviricotina* biology but rather a frequent occurrence in their evolutionary history.

**Figure 5.**
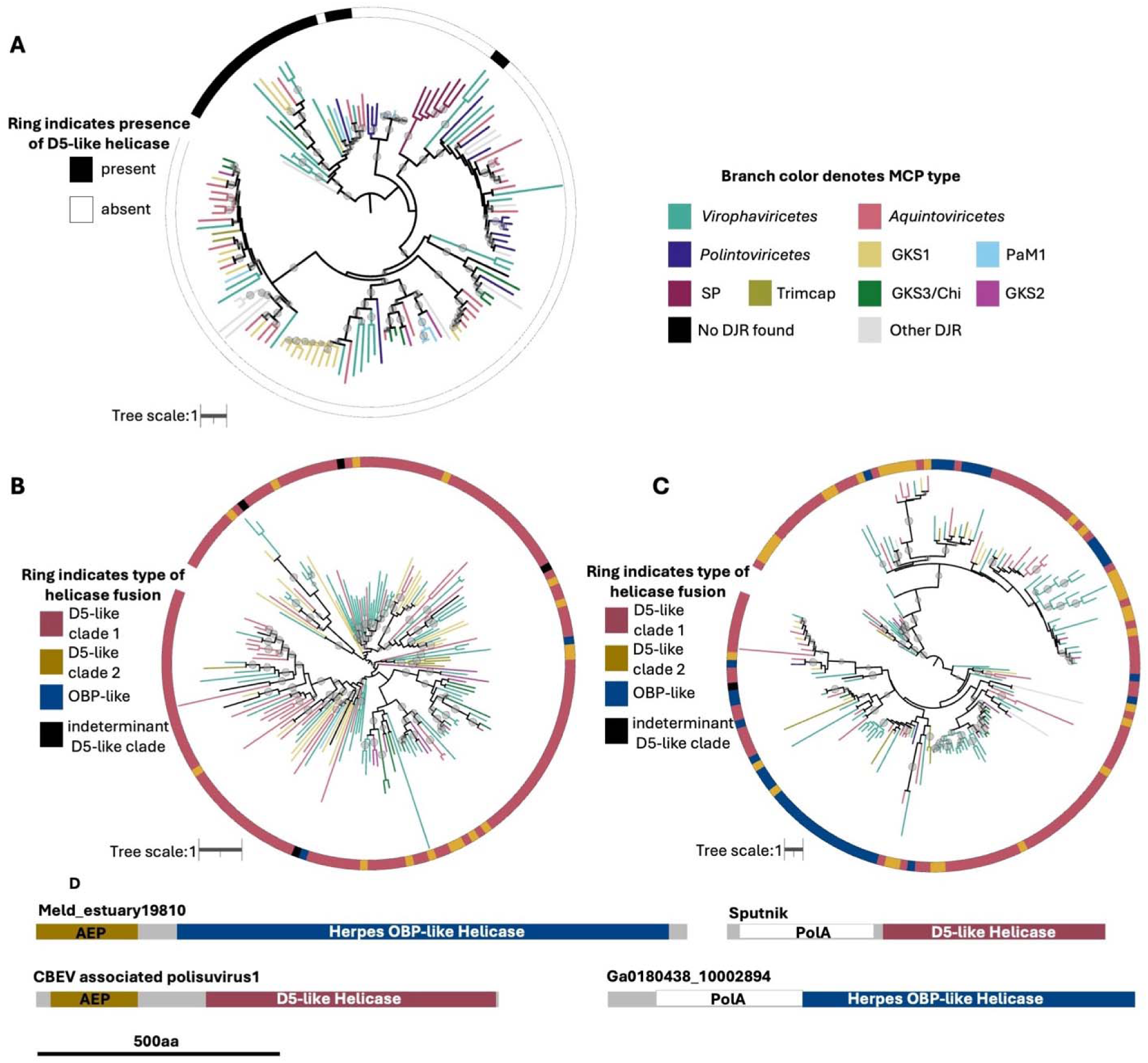
Polisuvirus replicases show whole gene and sub-genic mosaicism. Amino acid alignment based tree of *Polisuviricotina* encoded PolB proteins (A), PolA protein domains (B), or AEP protein domains (C). Branch color denotes MCP type. For (A) the outer ring denotes the presence of a stand-alone D5-like helicase. For (B) and (C) the outer ring denotes the type of C-terminal helicase domain fused to the PolA or AEP domain. D5-like clade 1 corresponds to OG002, while D5-like clade 2 corresponds to OG000_M. All trees are artificially rooted at the midpoint. Bootstrap values greater or equal to 90 are indicated as circles on branches. D. Examples of different replicase architectures found in polisuviruses.

### Domain level mosaicism of replicases

In addition to the genome-level mosaicism we found in these viruses, we also uncovered widespread protein-level mosaicism in the replicase proteins that they encode. Apart from pPolB, the most common replicase types in our genome set were PolA and AEP proteins. Typically, the PolA or AEP domains occupy the n-terminal portion of a replicase and are fused to a c-terminal D5-like helicase domain (Figure 5 B, C, and D). PolA and AEP fusions to superfamily 3 (SF3) helicase domains have been noted to occur in virophages and PLVs previously(Koonin and Krupovic 2017). The PolA-SF3 fusion is known to be particularly common, and proteins with this domain architecture have been dubbed ‘TVpols’(Iyer et al. 2008; Gupta et al. 2017). We find that AEP helicase fusions are also common amongst polisuviruses, with the majority of helicase domains belonging to a D5-like helicase and a sizable minority belonging to a Herpes OBP type helicase. We also detected diversity within the D5-like helicase domains which occurred as two separate orthologous gene groups (OG002 and OG000_M) in PolA and AEP proteins. Two additional OGs were annotated as D5-like helicases, OG203 and OG211, but these OGs were not particularly abundant, and did not occur as fusions with other recognizable domains (Table S1, Data S1).

We made amino acid alignment based trees for the PolA and AEP domains in our genome set to see if these proteins underwent horizontal transfer in a manner similar to the pPolB proteins. The tree revealed that PolA and AEP domain clades did not correlate well with MCP-type (Figure 5B and C). Additionally, we saw exchange of the helicase domain within PolA and AEP clades. This was particularly pronounced amongst AEP proteins which appeared to have frequent shuffling of the helicase domain. The PolA domain appeared to have a more stable relationship with the D5-like helicase clade 1 (OG002) but we did see some associations with D5-like helicase clade 2, and two instances of PolA-OBP fusions in our dereplicated genome set (Figure 5B). These results suggest that in addition to horizontal transfer of full replicase genes there is also swapping of individual replicase domains between polisuviruses, indicating that these viruses have mosaicism on the genome and gene level.

### Viral genomes reveal potential mechanisms for replicase mobility

The widespread mosaicism of *Polisuviricotina* replicases raises the question of what mechanisms may be responsible for replicase turnover. Hence, we examined individual genomes manually to uncover some insight into what mechanisms might contribute to the exchange of replicase proteins. Interestingly, we found that a number of genomes were found to encode replicases of multiple classes (Figure 6). Some of these genomes appeared to be variably degraded endogenous elements (Figure 6A,B, and D), but others appeared to be intact or nearly intact genomes (Figure 6C,F,G and H). In multiple cases, genomes were integrated in tandem (Figure 6 A,B, and E). While this phenomenon is less explored in eukaryotes, tandem integration of mobile elements has been observed in bacteria and has been proposed to contribute to MGE mosaicism by facilitating cycles of accretion and deletion of tandem sequence(Pavlovic et al. 2004; Bellanger et al. 2011). Deletions of internal sequence overlapping two integrated viruses could facilitate the reassociation of replicative and morphogenic modules. Notably, in some anemone genomes, we identified tandem integration of polinton elements and PLVs (Figure 4A and B). This is consistent with the previous observation that ‘polintons’ from the sea anemone *Nematostella vectensis* had a PLV-like MCP(C. Bellas et al. 2023).

**Figure 6.**
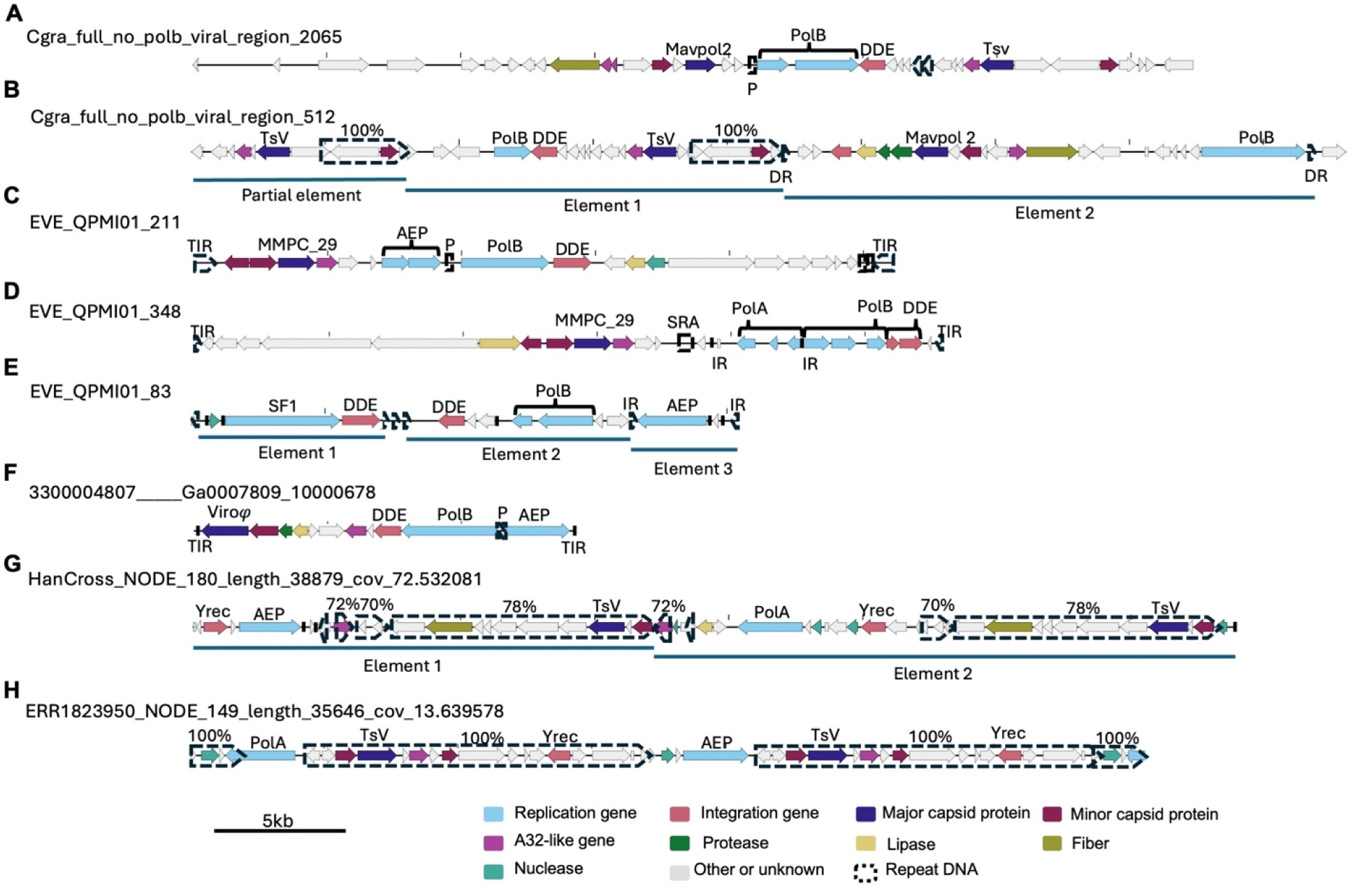
Polisuviruses display tandem integration and acquisition of additional replisomes. Diagrams of *Polisuviricotina* genomes. Genes are represented as solid arrows and their functions are denoted by color. Repeat sequences are represented by dashed arrows, with the level of shared nucleotide identity indicated above each arrow. Approximate boundaries of tandem elements are indicated where appropriate. TIR is Terminal inverted repeat. IR is inverted repeat. DR is direct repeat. SRA is small repeat array. P is palindromic repeat.

We also observed instances where a single element appeared to have acquired a second replicase (Figure 6C, D, and F), as well as two environmental metagenome derived contigs that possessed tandem mosaic copies of viral genomes (Figure 6G and H). The first of these contigs, HanCross_NODE_180_length_38879_cov_72.532081, consists of two homologous tandem morphogenic modules (70 to 78% nt identity) with conserved synteny but one copy is associated with a PolA gene and the other with an AEP gene (Figure 6G). The other contig, ERR1823950_NODE_149_length_35646_cov_13.639578 has nucleotide identical copies of morphogenic genes with only the replicases distinct between the two genomes (Figure 6F). These genomes appear to represent rare examples of recent replication turnover, and are consistent with the patterns of replicase mosaicism that we observe through phylogenetic analysis (Figure 2, Figure 5).

In multiple instances of genomes with multiple replicase classes, sets of repeats separate from the terminal inverted repeats (TIRS) bracketed one set of replication machinery apart from the morphogenic module (Figure 6C, D and E). This arrangement is consistent with mobilization of the replicases and suggests that these *Polisuviricotina* replicases may be capable of moonlighting as autonomous mobile elements. The concept of an independent replication-integration module has previously been established by the recognition of transpovirons, which possess pPolBs and DDE transposases but no recognized morphogenic module(Koonin and Krupovic 2017). Some of the multi-replicase genomes appear consistent with mobilization of a transpoviron into a *Polisuviricotina* genome (Figure 6C) while in others it looks like a PolA or AEP gene has been acquired (Figure 6D and F). One contig, EVE_QPMI01_83 appears to be three elements with two resembling transpovirons and the third consisting solely of an AEP gene (Figure 6E). Interestingly, while the pPolB genes in these examples are consistently paired with DDE integrases, the PolA and AEP genes are not always associated with integration genes, suggesting that they may be able to mobilize independently. Some of these replicases are encoded near endonuclease genes (Figure 6C, E, G and H). Homing endonuclease genes (HEGs) are known to facilitate replicase gene conversion in bacteriophages(Goodrich-Blair and Shub 1996), and putative HEGs are also associated with PolB genes in eukaryote infecting nucleocytoviruses(Deeg et al. 2018; Erazo-Garcia et al. 2025). HEGs could conceivably mobilize PolA or AEP genes through a collaborative homing process similar to what has been observed for HEGs and neighboring self-splicing introns. In this hypothetical scenario, polymerase-primase genes could aid HEG mobility by copying the primase and HEG locus into a HEG generated cleavage site.

Transpovirons are known to sometimes integrate into nucleocytovirus genomes(Desnues et al. 2012). We reasoned that if *Polisuviricotina-*encoded PolA and AEP replicases are capable of autonomous mobility, then they would likely mobilize into some nucleocytovirus genomes. One of the core genes of nucleocytoviruses, the primase-polymerase D5, consists of an N-terminal AEP domain and a C-terminal D5-like SF3 helicase domain similar to many polisuvirus replicases. We reasoned that additional D5 homologs in the genomes of nucleocytoviruses might be mobile genes rather than duplications of the core D5 gene. A remarkable case of this occurs in *Theiavirus salishense*: the Bodo Saltans virus (BsV) strain NG1(Deeg et al. 2018) which encodes 12 AEP-helicase fusions throughout its genome (Figure 7A). 11 of these genes showed a close relationship, while the last one appeared distinct (Figure 7). An amino acid alignment based tree of the AEP domains with *Polisuviricotina* AEP proteins and nucleocytovirus D5 proteins showed that the unique AEP-helicase fusion in BsV clustered with nucleocytovirus D5 homologs while the others clustered with *Polisuviricotina* proteins (Figure 7B) suggesting that the additional BsV AEP-helicase fusions may be mobile elements related to polisuvirus replicases. Additionally, the tree suggested occasional acquisition of D5 homologs from nucleocytoviruses by polisuviruses, and occasional transfer of polisuvirus-encoded replicase homologs into other nucleocytovirus genomes (Figure 7B). The additional AEP-helicase homologs in BsV lack signatures of mobility, such as flanking repeats; however, some mobile elements, such as HEGs, also lack these signatures, and flanking accessory sequences would not necessarily be needed for a primase-polymerase to template its own replication into broken DNA. AEP proteins are functionally versatile, with some being capable of primase, terminal transferase, translesion synthesis, and processive polymerase activity (Guilliam et al. 2015). How this versatility may have impacted MGE evolution warrants further investigation.

**Figure 7.**
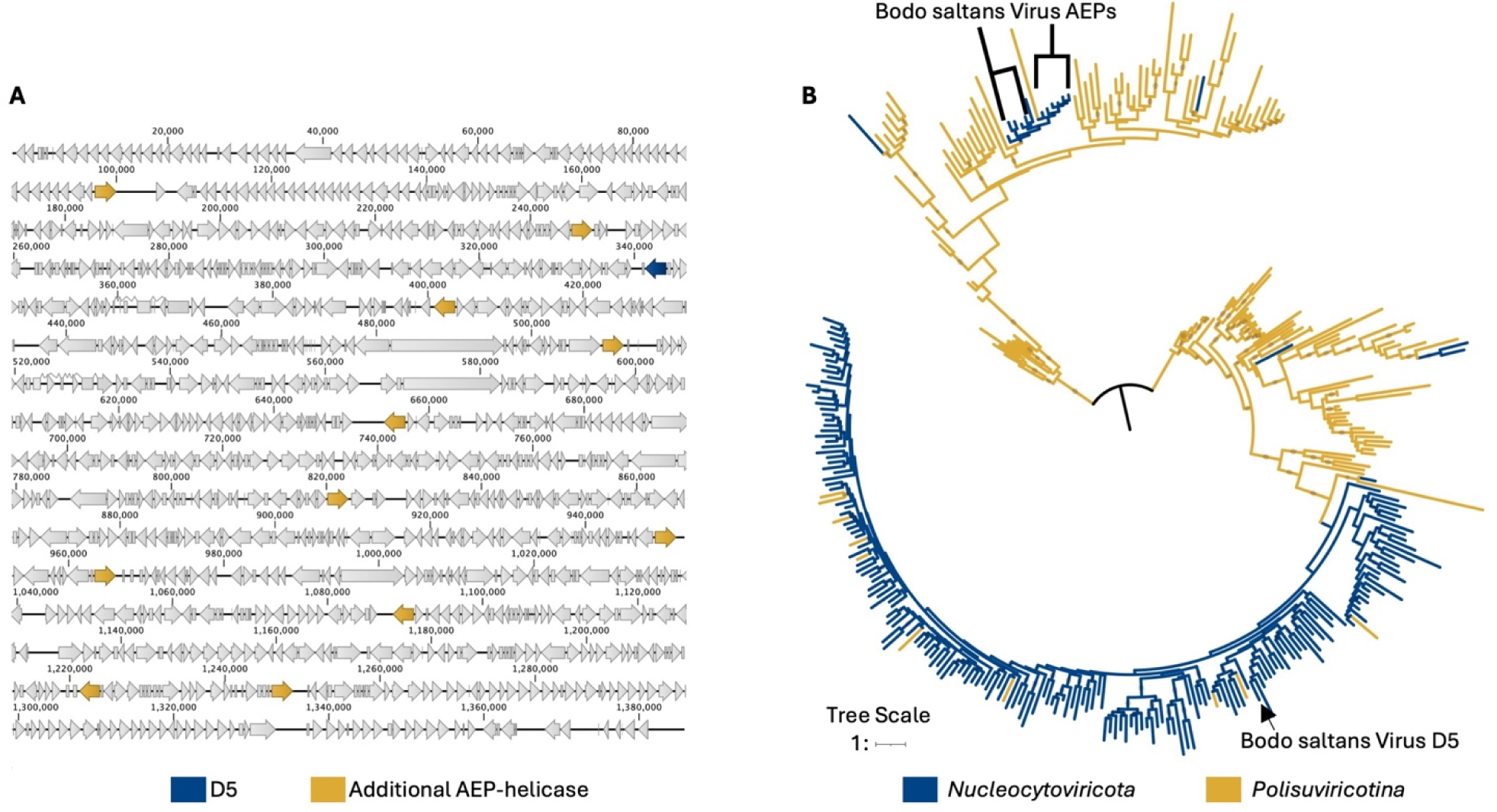
Distribution of AEP-SF3 genes in BsV suggests gene mobility. (A) A diagram of a Bodo saltans virus strain NG1 (NC_075036). Genes are represented by arrows. A blue arrow denotes the D5 primase-polymerase. Yellow arrows denote additional AEP-helicase fusion genes within the NG1 genome. (B) An amino acid alignment based tree of AEP domains from both *Polisuviricotina* and *Nucelocytoviricota*. The classification of the AEP encoding genome is denoted by branch color. The tree was artificially rooted at the midpoint. Bootstrap values greater than 90 are indicated by circles on branches.

### Conclusions

Here we show that extensive mosaiscism has been a feature of polisuvirus evolution, with multiple lineages of morphogenic modules cross associating with different lineages of replication machinery. This mosaicism poses a challenge for polisuvirus taxonomy; we propose that the evolutionary history of the morphogenic module should take priority over that of the replication module when classifying polisuviriuses. Classification on the basis of morphogenic genes is already the case for several higher level viral taxa(Koonin et al. 2020; Koonin, Kuhn, et al. 2024) and would be mostly concordant with our phylogenetic results (Figure 1). Evolutionary history of the replication module may be informative for classification at lower taxa levels but hybrid lineages are likely to continue to emerge.

It remains unclear why *Polisuviricotina* replication modules turnover so frequently. Our examination of individual *Polisuviricotina* genomes with multiple replicases raises the possibility some *Polisuviricotina* replication modules may have autonomous mobility. This hypothesis remains speculative and preliminary as extensive experimental work would be needed to antiviral systems of their hosts. This has been suggested for hyperparasites of bacteriophages which exhibit replisome mosaicism where the replisome is a known target for bacteriophage-encoded anti-parasite defenses(Barth et al. 2021, 2023). We have yet to observe cellular or giant virus defenses that specifically target *Polisuviricotina* replication modules, but the presence of many genes with putative defense functions in *Polisuviricotina* genomes Figure 1, suggests that they are likely engaged in a co-evolutionary arms race with cellular hosts, viral hosts, or other small viruses.

The unstable association of *Polisuviricotina* morphogenic and replicative modules draws an interesting parallel to small MGEs in bacteria which frequently exhibit mosaicism(Pesesky et al. 2019; Wozniak and Waldor 2010; Benler et al. 2021; Dion et al. 2020; Hendrix et al. 1999). A possible explanation for the mosaicism of very small genomes is the complexity hypothesis. The complexity hypothesis states network connectivity determines the transferability of genes between genomes(Cohen et al. 2011; Jain et al. 1999). If replication functions are not functionally integrated into the other functions of an MGE or virus, such as morphogenesis, then replication functions are capable are re-associating with other mobile genomes. The same principle could also explain why MGE cargo genes, like antimicrobial resistance genes and anti-viral defense systems, are so frequently exchanged. The replisomes for larger genomes, such as cells and nucleocytoviruses, are more complex and involve more interacting proteins. This greater complexity likely acts as a barrier to horizontal gene transfer and further integrates the replisome into the evolutionary history of the whole genome.

The highly mosaic nature of *Polisuviricotina* genomes has important implications for understanding the biology and evolution of these elements. In their simplest forms, *Polisuviricotina* genomes appear to be consortiums of morphogenic and replicative modules. The unstable associations of these modules suggest that each are able to evolve independently of each other. In the case of replicative modules, the existence of transpovirons and mobile AEP-helicase insertion sequences suggests that some modules may be able to function as their own autonomous mobile elements, adding an additional layer of complexity to the web of antagonistic interactions between eukaryotes, nucleocytoviruses, and polisuviruses. The evolutionary entanglement of these entities has likely set the stage for the staggering diversity of *Polisuviricotina* by providing ample opportunities for horizontal gene transfer and the strong selective pressures of a multi-level coevolutionary arms race. Our findings illustrate how rampant mosaicism underpins the diversity of small DNA viruses.

## Materials and Methods

### Generation and curation of a polisuvirus database

Previously published polisuvirus genomes from several studies(Yutin, Shevchenko, et al. 2015; Bao et al. 2015; Paez-Espino et al. 2019; C. M. Bellas and Sommaruga 2021; Jeong et al. 2023; C. Bellas et al. 2023; Starrett et al. 2021; Buck et al. 2024; Stephens et al. 2024; Koslová et al. 2024), were collated into a single set of 10,246 genomes. Initial dereplication was done at 95% identity using dRep (dRep dereplicate -l 1500 -pa 0.9 -cm total --warn_aln 0.8 -p 32). This resulted in 3504 genomes, which was still too large for phylogenetic analysis, and so a subsequent round of dereplication was performed with hclust leaving a final set of 1055 genomes. To dereplicate this database, orthofinder was run on a subset of the database (1273 genomes), to generate orthogroups (OG), and then the hclust function in R was used to dereplicate the database on the basis of OG occurrence.

### Generating gene content profiles

Orthofinder(Emms et al. 2026) was ran on the dereplicated database of polisuvirus genomes to generate a new set of OGs. Manual curation of this second set of OGs, in particular splitting of certain OGs into separate subclusters was performed to generate the final set of OGs. To identify subclusters, all proteins in an orthologus group were compared to each other using blastp (e-value 1e-3), and results were subsequently visualized in igraph. In these OGs, only the largest subcluster that could be visualized in igraph was retained. For each curated OG, the constituent protein sequences were aligned using MUSCLE5 with default parameters(Edgar 2022). Alignments were trimmed using TrimAl and default parameters(Capella-Gutiérrez et al. 2009). The Hmmbuild function from HMMER3(Eddy 2011) was used to produce HMM profiles for each trimmed alignment.

In a few cases, alignments revealed certain OGs to be comprised of highly mosaic protein sequences. This was particular noticeable for OGs containing D5-like or OBP-like helicase domains fused to either PolA or AEP domains. Manual sequence trimming and realignment of trimmed sequences was used to generate alignments specific to helicase or polymerase-primase functional domains, and these curated alignments were used to generate HMMs for our final set. HMMsearch function from HMMER3 was used to generate gene profiles for each genome. These gene profiles were then used to generate binary files for each genome.

We modified the virtree algorithm to account for multiple hits. The revised Virtree code can be found at https://github.com/faylward/viralrecall. To account for subgenic mosaicism, we allowed polisuvirus proteins to have multiple hits to different OGs and treated each unique OG combination as distinct trait for the purpose of the binary fasta profiles.

### OG annotation

Functions for OGs were predicted using the HHPRED tool from the Max Planck Institute (MPI) Bioinformatic Toolkit(Zimmermann et al. 2018; Gabler et al. 2020). For each OG, a multiple sequence alignment of the constituent protein sequences as well as three individual protein sequences were used as a query against the PDB_mmCIF70, Pfam-A, UniProt-SwissProt-viral70, and NCBI_ConservedDomains(CD) databases.

For assigning MCP OGs to recognized MCP clusters, the individual proteins within the OGs were cross referenced with previously published MCP clusters(C. M. Bellas and Sommaruga 2021; C. Bellas et al. 2023).

### OG distribution analysis and alluvial plots

Alluvial plots were generated using the geom_alluvium command in the R package ggalluvial(Brunson 2020). Phylogenies of polisuvrius genome binary profiles were generated from the binary fasta file using IQTREE (-st BIN -m GTR2+ASC -nt AUTO -bb 1000 -wbt -seed 123 --runs 10). For single gene and subgenic domain trees, sequences were aligned with MUSCLE5, and trimmed with TrimAl(Edgar 2022; Capella-Gutiérrez et al. 2009). Phylogenies were generated from the trimmed alignments using IQTREE (-m LG+F+R10 -bb 1000 --wbtl -T AUTO --runs 10).

Phylogenetic trees were visualized using iTol(Letunic and Bork 2024).

## Data Availability

Genomes, HMM profiles, alignments, and Newick format tree files used for this study are available at https://doi.org/10.5281/zenodo.23740701.

## Acknowledgements

We would like to thank members of the Aylward lab for useful discussion.

---

**Figure S1.**
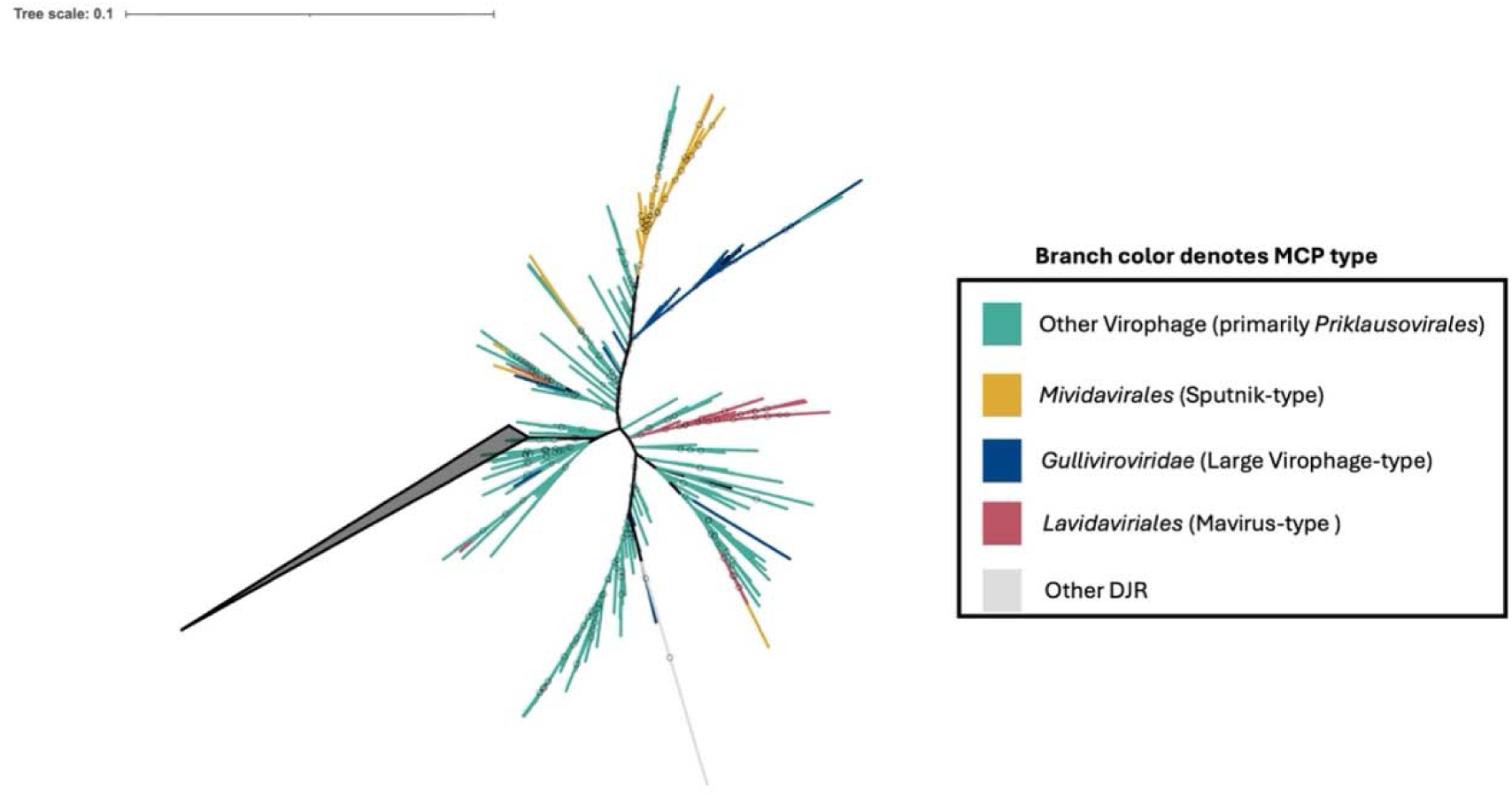
A different view of the same tree from Figure 1, with the non-*Virophaviricetes* clades collapsed.

**Figure S2.**
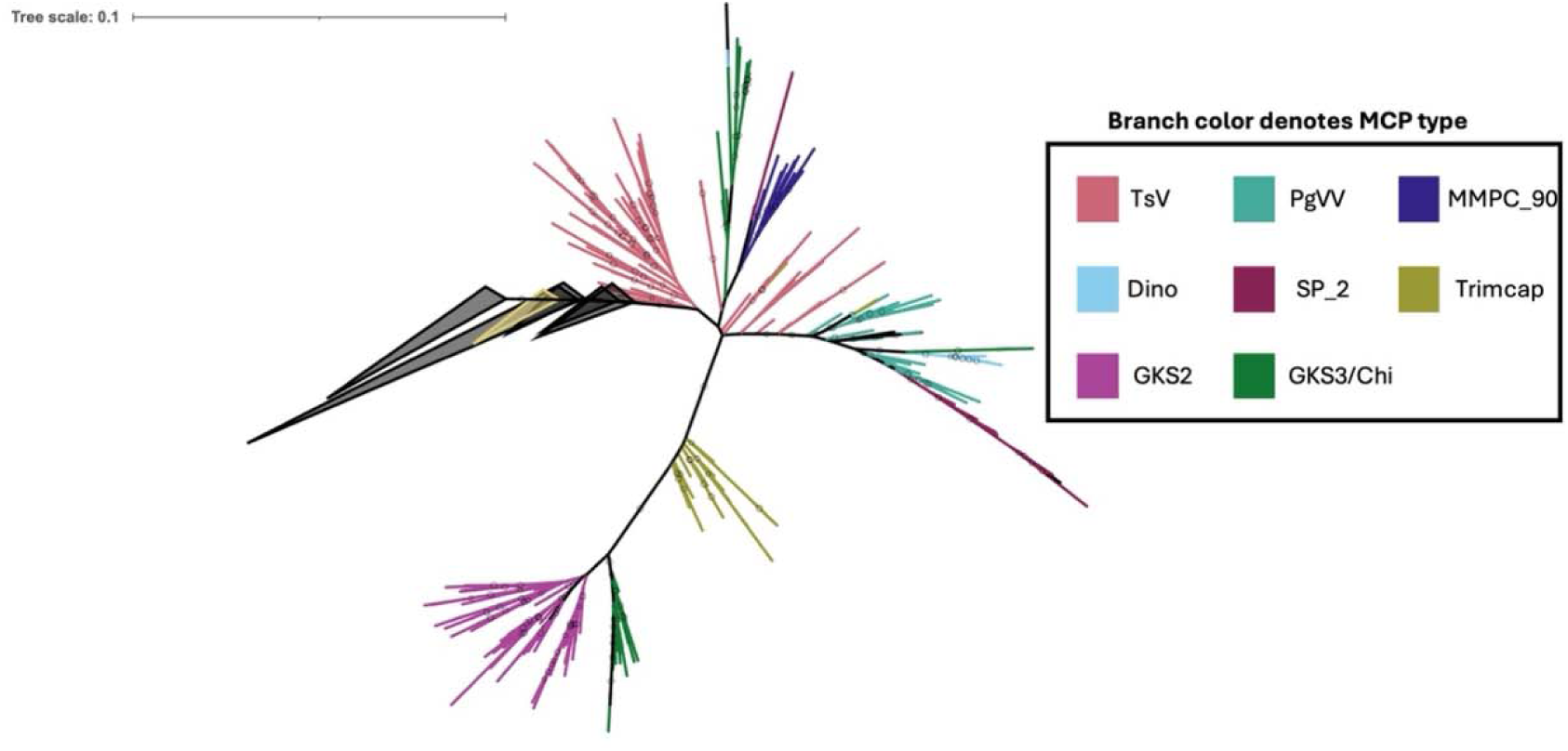
A different view of the same tree from Figure 1, with the *Virophaviricetes, Polintoviricetes,* and GKS1 clades collapsed.

